# Comparative Assessment of Large Language Models for Microbial Phenotype Annotation

**DOI:** 10.1101/2025.11.24.690272

**Authors:** Philipp C. Münch, Nasim Safaei, René Mreches, Martin Binder, Yichen Han, Gary Robertson, Eric A. Franzosa, Curtis Huttenhower, Alice C. McHardy

## Abstract

Large language models (LLMs) are increasingly used to extract knowledge from text, yet their coverage and reliability in biology remain unclear. Microbial phenotypes are especially important to assess, as comprehensive data remain sparse except for well-studied organisms and they underpin our understanding of microbial characteristics, functional roles, and applications. Here, we systematically assessed the biological knowledge encoded in publicly available LLMs for structured phenotype annotation of microbial species. We evaluated the performance of over 50 LLMs, including state-of-the-art models such as Claude Sonnet 4 and the GPT-5 family of models. Across phenotypes, LLMs reached accurate assignments for many species, but performance varied widely by model and trait, and no single model dominated. Model self-reported confidence is informative, with higher confidence aligning with higher accuracy, and can be used to prioritize phenotype assignment, effectively distinguishing between high-and low-confidence inferences. Overall, our study outlines the utility and limitations of text-based LLMs for phenotype characterization in microbiology.

## Introduction

The exponential growth in scientific publications has led to a wealth of knowledge dispersed across numerous databases and platforms (Reimer et al. 2019; Meier-Kolthoff et al. 2022; Whitman 2015). However, much of this is provided as unstructured text and paired with sparse, heterogenous, or missing metadata, complicating discovery and integration. The diverse and distributed nature of these resources poses substantial challenges for researchers seeking to comprehensively understand a topic (Brbić et al. 2016). This also limits reuse in machine-learning-based efforts, which often require high-quality FAIR data available in structured, machine-actionable formats (Wilkinson et al. 2016).

Large language models (LLMs) are increasingly used for large-scale text extraction. These models are typically trained on vast and largely unspecified text corpora from the internet (Buck et al. 2014; Brown et al. 2020), which contain billions of web pages across diverse domains and languages. Consequently, the resulting models capture and generalize from an extensive cross-section of human knowledge, including both everyday discourse and specialized scientific expertise. This broad training also encompasses substantial amounts of scientific literature, such as research articles, preprints, and technical documentation that have been made publicly available online through open access repositories, institutional websites, and scientific databases.

Microbial phenotype data describe the traits of microorganisms, such as morphology, metabolism, growth conditions, and ecological roles that result from the interaction between their genetic makeup and environment. These data can be found in diverse sources, such as primary research literature, taxonomic monographs like Bergey’s Manual (Whitman 2015), and curated databases such as BacDive (Schober et al. 2025) and the List of Prokaryotic Names with Standing in Nomenclature (LPSN)(Parte 2018). Typically, phenotypic information is compiled through expert curation, where researchers extract data from the literature and laboratory records to create structured datasets. These resources capture only a slice of the available phenotypic information due to the time-consuming and labor-intensive nature of manual curation. Achieving comprehensive coverage and timely incorporation of new information with this approach remains challenging. LLMs offer a promising complement to this approach due to their capabilities for rapidly extracting and synthesizing information from text, and to create structured, machine-actionable FAIR data records.

In this study, we evaluated the capacities of available large language models to query and integrate microbial phenotypic data from multiple sources (Dutilh et al. 2013). Specifically, our study addresses three questions: (i) how accurate and consistently can LLMs assign microbial phenotypes to microbial taxa (ii) to what extent do they recognize and quantify their own uncertainty, and (iii) how can hallucinations, which are one of the most pervasive challenges in LLM-based knowledge extraction, be identified and mitigated. To facilitate future model comparisons and assessments, we provide a comprehensive benchmarking framework and high-quality, structured, and machine-readable microbial phenotype data.

## Results

### Platform for reproducible benchmarking of general-purpose LLMs

To assess the microbial knowledge captured by large language models (LLMs), we established a benchmarking setup to evaluate general-purpose models on domain-specific microbial knowledge tasks under controlled, reproducible conditions. Our evaluation was conducted in a zero-shot setting, where models were queried directly without any task-specific fine-tuning, contextual examples, retrieval augmentation, or access to external databases. The platform ensures comparability across models through standardized species sets and phenotype metrics, enabling consistent evaluation of knowledge depth, accuracy, and calibration. Beyond assessing individual models, it provides a scalable reference for tracking progress in biological reasoning across successive LLM generations.

To implement this approach, we developed a browser-accessible framework specifically designed to automate the end-to-end evaluation of LLMs for microbial phenotype inference tasks. The system accepts a list of biological entities (e.g., species names) and a set of query and validation templates, then automates standardized prompting, response parsing, and comparison to ground-truth datasets (**Supplementary Fig. 1**). The framework is model-agnostic and supports multiple LLM providers through OpenRouter, an API aggregation service that ensures consistent and reproducible access to a diverse set of LLMs. The framework incorporates several key features for robust evaluation. It uses a machine-readable output schema for harmonized evaluation across experiments and incorporates mechanisms for output validation and error handling, such as retries for malformed responses or API failures. Unified prompt templates enable reproducible experimentation across models and traits, while built-in logging ensures traceability of all evaluations. While this framework can be extended to related microbiology tasks, such as growth conditions or metabolic capabilities, we focused here on microbial phenotypes as a well-defined and diverse demonstration case.

We make all results available through a browser-based dashboard that renders analyses directly from the evaluation database. As runs complete, the dashboard updates summary statistics and comparative plots in real time, allowing model- and species-set–level stratification without additional processing. Users can filter by model/provider and compare successive model generations. Visualizations are generated from the same machine-readable schema used for scoring, ensuring versioned, reproducible figures and exportable data.

Within this framework, we assembled a panel of over 50 large language models to capture the current diversity of the LLM landscape. The panel spans systems released between 2023 and 2025, encompassing multiple model families, scales, and both closed-source and open-source architectures. It includes proprietary systems (e.g., GPT-family, Claude, Gemini), widely available open-access models (e.g., DeepSeek, Llama, Gemma), and community-released options (e.g., Mixtral), covering both base and instruction-tuned variants. This diversity enables comparisons across architectures, parameter scales, and training philosophies, from compact models (<10B parameters) designed for local inference to frontier systems deployed via cloud APIs. We designed this panel to represent the spectrum of tools most likely to be used in research and applied microbiology, balancing readily accessible and lower-cost options with state-of-the-art commercial models such as GPT-5. This breadth also allows us to examine how model size, accessibility, and provenance influence the retention and calibration of microbial phenotype knowledge.

### Dataset for microbial phenotype inference

As the gold standard for binary and multiclass phenotypic traits of microbial species, we used bugphyzz, a comprehensive, curated phenotype dataset (**Fig. 1A, Supplementary Table 1**), including cell shape, sporulation, environmental associations, and pathogenicity traits. While primarily bacterial, the dataset also includes archaeal microbial species, thus covering multiple domains. For benchmarking, we stratified bugphyzz into two subsets: a well-annotated (WA) set (n=3,876) with no more than five missing trait values per species, and a less-annotated (LA) set (n=15,256; **Supplementary Dataset 1, 2, 3**). The former served as our main evaluation set, offering both data completeness and taxonomic breadth, as its phylum-level composition closely mirrors that of the broader dataset (**Fig. 1B**). This dataset includes pathogenicity-related targets, such as animal and plant pathogenicity, allowing us to evaluate each model’s performance in predicting potentially sensitive traits with implications for dual-use applications in microbiology.

**Figure 1:**
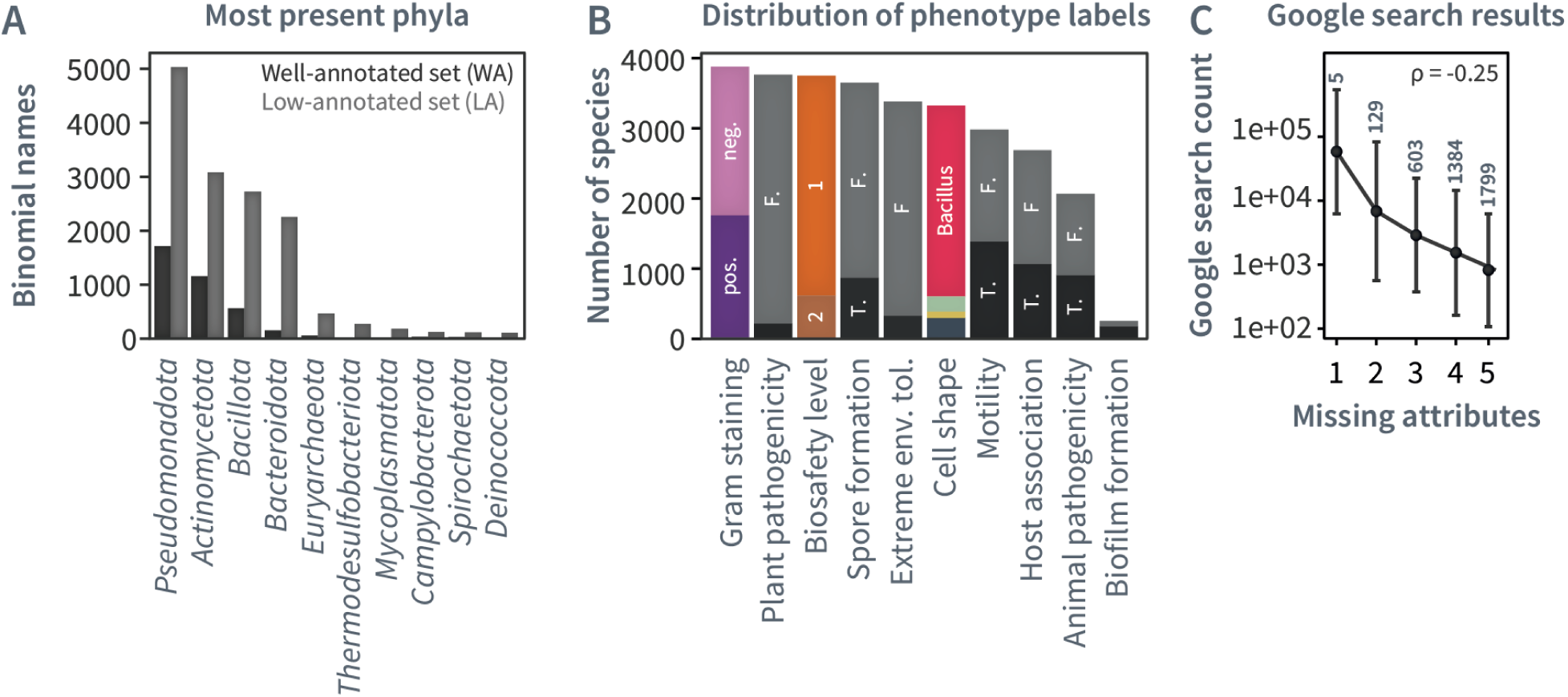
Overview of dataset characteristics. **A**) Distribution of the ten most prevalent phyla in the WA and LA subsets. **B**) Distribution of labels and annotation completeness across phenotypes in the WA subset. Abbreviations: pos./neg.: Gram+/Gram-; T.: TRUE; F: FALSE, 2: BSL category 2. **C**) Correlation between Google search results and annotation completeness. Species with fewer Google search results have more missing phenotypic attributes in the WA set.

To assess and optimize LLM performance for microbial phenotype prediction, we implemented a two-tier evaluation strategy. In the first phase, we conducted a comprehensive model comparison using the WA subset, as its richer annotations provide a stronger basis for validation. We processed all binomial names across different LLM models, enabling a thorough comparative analysis of their performance and identification of high-performing models for our tasks. In the second phase, we extended predictions to the more poorly characterized taxa in the LA subset, where sparse annotation makes direct validation less reliable. Based on the WA evaluation, we selected grok-4-mini as the best-performing model for systematic assessment of the LA subset, balancing predictive accuracy with computational cost.

Since LLMs are trained mostly on data derived from publicly accessible internet sources, we quantified how often binomial species names appeared as exact matches in Google Search results, as a proxy for training data coverage of the analyzed microbial taxa (**Supplementary Dataset 4**). The number of missing attributes was negatively correlated with Google hit counts (Spearman’s ρ=-0.25, p=2*10^-16^, one-sided), indicating that species with greater web visibility are both better annotated in bugphyzz and more accurately predicted by LLMs (**Fig. 1C**). This relationship shows that Google visibility, as a rough proxy for public data availability, aligns with both annotation completeness and predictive performance.

### Quality, consistency and hallucination-content of LLM-generated output

LLMs can generate convincing-sounding answers even when their knowledge of a topic is limited or nonexistent (Banerjee et al. 2024). This tendency to “hallucinate” poses a substantial challenge in validating the reliability of LLM-generated information, particularly in specialized fields such as microbiology. It is critical to avoid incorporating confidently asserted yet incorrect information into microbial research databases, which could potentially misdirect research efforts or lead to erroneous conclusions. To evaluate this issue, we established a method that leverages an LLM’s ability to self-assess its own knowledge base, serving as an effective filter against potential hallucinations.

The filtering method evaluates LLMs’ ability to self-rate their knowledge about specific microbial species and provide reliable phenotypic description. To this end, bacterial species are categorized into four knowledge classes: non-assigned (‘NA’), ‘Limited,’ ‘Moderate,’ and ‘Extensive’. We then designed three increasingly verbose queries (“Templates 1-3”) to assess this confidence level based on the binomial name (**Supplementary Dataset 5**). For example, query template 1 defines classes such as “Minimal to basic information available, challenging to make accurate predictions; ‘Moderate’: Moderate information available, including some phenotypic, morphological, genetic, or physiological characteristics […]”. Query template 3 adopts a more data-driven classification approach: “[…] Limited: Species with minimal data and research, typically with few species or subspecies (<5 species, <2 subspecies), little genetic information (<10 scientific articles), no complete genome sequences, and limited presence in culture collections (absent or very few species) […]”. Furthermore, template queries 2 and 3 contain the phrase “If the species name is not a real or recognized bacterial species, or if there is no information available to determine the knowledge level, respond with *NA*” to assess the model’s capability to recognize knowledge gaps and avoid hallucination.

To assess the robustness of the LLM’s confidence scoring system, we generated sets of artificial binomial names (n=50 each) across four categories (Fig. 2A, **Supplementary Dataset 6**). These included (i) completely artificial names consisting of combinations of English words (e.g., “Amber Field”); (ii) Latin words (e.g., “Solispira lumina”) and (iii) hybrid names, representing combinations of Latin words with real genus or species elements (e.g., “Mycobacterium ferrum”, “Temporibacter coli”). We then evaluated all LLM models using these 200 artificial binomial names, applying the most comprehensive query template 3 to assess each model’s ability to correctly identify and categorize non-existent species (Fig. 2B, **Supplementary Dataset 7**). For larger LLMs such as OpenAI GPT-4 or Anthropic’s Claude 3.5 Haiku, the proportion of misclassified terms for each query type is low (with 0%, 0.5%, and 1.5% misclassified artificial binomial names for high, medium, and low verbosity, respectively). In contrast, the smaller and more cost-effective model (llama-3-8b-instruct) showed reduced misclassification rates with increasing query verbosity (misclassified terms: 98%, 97.5, and 74%, for low, medium, high verbosity, respectively). This suggests a potential interaction between model size and the benefit derived from more verbose queries.

**Figure 2:**
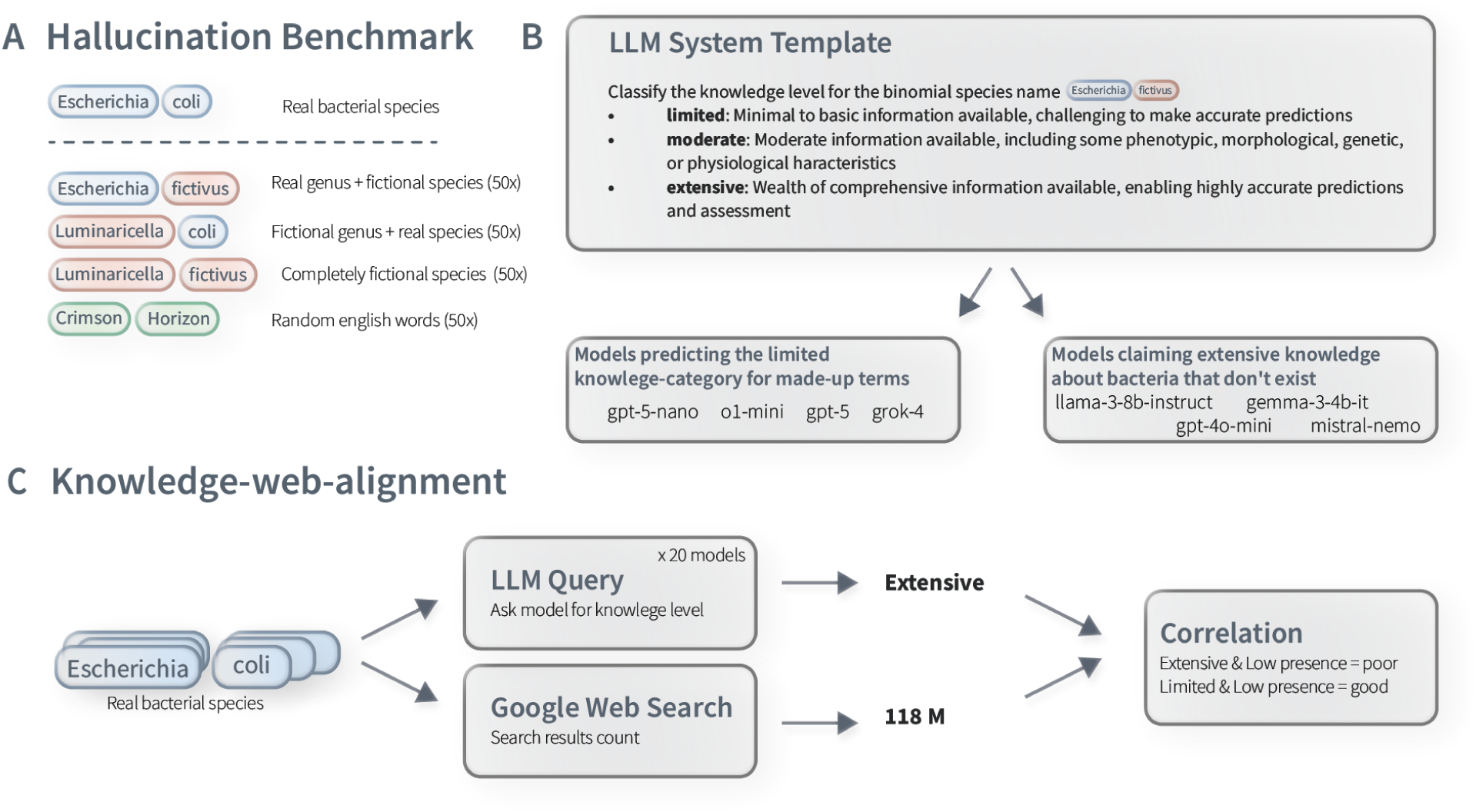
Overview of the LLM-based microbial phenotype evaluation framework. **A**) Schematic of synthetic binomial name generation for hallucination assessment. Four categories of fictional species names were created. **B**) Each synthetic name was evaluated using prompts that asked LLMs to self-rate their knowledge level. **C**) Framework for validating self-rated knowledge against web information availability. For each real binomial species name, models provided knowledge ratings, which we compared against the number of Google search results for that species.

We then evaluated 57 large language models on the hallucination benchmark data using the more verbose query template 3, which explicitly allows models to respond with “NA” for unknown species. From 11,400 model inferences overall, 66% (7,526) correctly generated NA rejections and 0.3% (38) inference failures/no-results and 33.6% (3,835) hallucinations, where models incorrectly provided a knowledge group assignment for non-existing species (**Supplementary Dataset 7**). We observed substantial variations in hallucination rates from 0.5% to 92% for different models. The top-performing model, gpt-5-nano, achieved 99.5% accuracy in correctly identifying artificial taxa as unknown. Other high-performing models include gpt-5 (97.5% accuracy) and grok-4 (96.0% accuracy, **Supplementary Dataset 8**). Conversely, we identified models with significantly higher hallucination rates. The worst performer, llama-3.2-3b-instruct, correctly groups 8% of artificial taxa in the NA/unknown group. These results highlight a substantial variability in model reliability when handling fabricated microbial taxa and underscore the importance of careful model selection for microbiological applications.

### Knowledge calibration and model agreement on real microbial species

We then applied this knowledge rating strategy to the well-annotated (WA) dataset to evaluate the LLMs (**Supplementary Dataset 9**). Since there is no ground truth for the correct knowledge group assignments, as appropriate classification depends on each model’s training data, which in most cases are not publicly accessible, we employed the Shannon entropy of model assignment consistency, a measure quantifying how consistently models assign each species to the same knowledge group, as an indirect measure of classification uncertainty. We complemented this with an alignment analysis using Google search counts to assess whether model classification correlates with real-world information availability (Fig. 2C). Knowledge group distribution varied substantially across the evaluated models (Fig. 3A). Of the 3,876 species in the WA set, only 91 species achieved consensus across all models, while for the majority models disagreed: 1,046 species were assigned to two knowledge groups, 2,291 species across three groups, and 456 species across all four different knowledge categories (including ‘NA’) (Fig. 3B, **Supplementary Dataset 10)**. Species with the highest classification uncertainty (entropy) included members of *Streptomyces* and species such as *Metamycoplasma arthritidis* (Fig. 3C).

**Figure 3:**
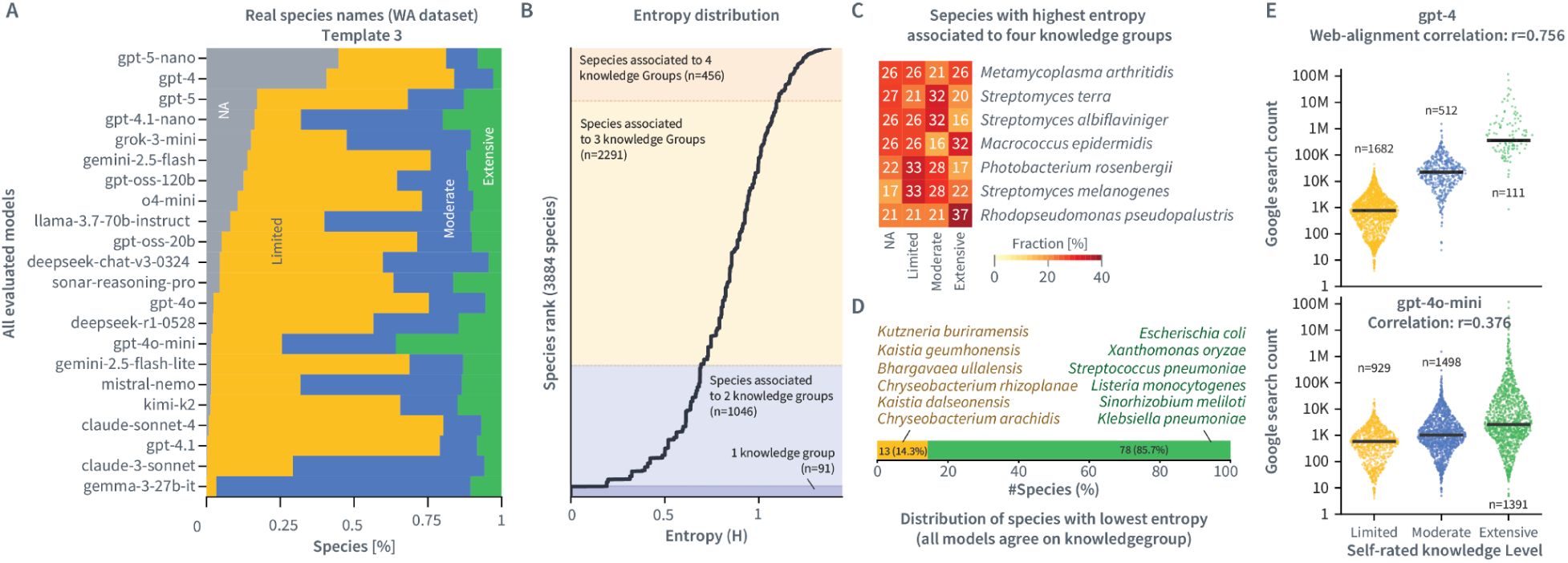
Inter-model agreement on knowledge group classifications. **A**) Distribution of knowledge group assignments across evaluated models and species in the WA dataset. **B**) Species ranked by classification uncertainty (Shannon entropy) using query template 3. Each point represents one species, sorted from highest disagreement (top) to highest agreement (bottom). Low-entropy species at the bottom reflect unanimous classification, where all or nearly all models assigned the same knowledge group. Colored regions indicate entropy thresholds corresponding to species split across two, three, or four different groups. Species in the upper region show the greatest inter-model disagreement, with models distributing classifications across multiple knowledge categories (see panel C for examples). **C**) Heatmap showing the seven species with the highest classification entropy from panel B, illustrating how different models split their knowledge group assignments for these ambiguous cases. Each row represents one model, and colors indicate the assigned knowledge group (percent). **D**) Distribution of knowledge-groups for the 91 taxa achieving full consensus (entropy = 0) across all tested models, categorized by their unanimously assigned knowledge group. Well-characterized species like *E. coli* were consistently classified as ‘Extensive’ (green), while recently described species like *Chryseobacterium arachidis* were classified as ‘Limited’ (yellow). **E**) Relationship between knowledge group assignment and web information availability. Distribution of Google search hit counts for microbial species in each knowledge category. Different models assigned varying numbers of species to each group, resulting in model-specific correlations between self-assessed knowledge level and web presence.

In contrast, species with strong inter-models agreement reflected expected patterns based on research prominence. Well-characterized species like *E. coli* and *Klebsiella pneumoniae* were consistently classified in the ‘Extensive’ knowledge group (Fig. 3D), while in the ‘Limited’ group contained recently discovered species such as *Chryseobacterium endophyticum,* reported in 2017 (Lin et al. 2017) and *C. ginsengisoli,* reported in 2013 (Nguyen et al. 2013). This differentiation between well-established and recently described species validates our knowledge classification system and demonstrates its ability to reflect the current state of microbial research. The approach provides a reliable metric for assessing the extent and accuracy of knowledge contained within LLMs about specific microbial taxa.

To evaluate how well model classifications align with real-world knowledge availability, we compared their knowledge grouping with Google search result counts retrieved using the exact binomial species name. The taxa with the largest number of indexed pages were, as expected, *Escherichia coli* (118M hits), *Staphylococcus aureus* (70M), and *Pseudomonas aeruginosa* (34M). In contrast, lesser-known species such as *Qipengyuania xiamenensis* and *Qipengyuania sphaerica* returned 4 and 5 hits, respectively. We then calculated a web alignment score for each model by correlating its knowledge-level assignments with the log-transformed Google hit count. Models varied in their degree of alignment; gpt-4 showed a strong correlation (0.756), while gpt-4o-mini had a weaker correlation (0.376), largely because gpt-4o-mini classified far more species into the ‘Extensive’ knowledge group (n=1,391) than gpt-4 (n=111) (Fig. 3E). Further, gpt-4o-mini classified 1,391 species as having ‘Extensive’ knowledge, compared with only 111 for gpt-4, suggesting that smaller instruction-tuned variants may overestimate their familiarity with biological entities or use different internal thresholds for knowledge assessment. Models such as gemma-3-27B, and certain earlier Claude variants showed poor alignment, likely due to differences in training data composition. Interestingly, gpt-5-nano and gpt-4 also assigned around twice as many species to the “NA” category compared with other models, suggesting that their higher correlation score stemmed in part from more conservative, less overconfident classifications.

### Benchmark reveals LLM proficiency and limitations in phenotype prediction

Having established each LLM’s ability to effectively self-assess its knowledge about species, we next evaluated their performance in assigning phenotype characteristics to microbial species. We applied 31 LLMs (selected based on usage costs) to all binomial species in the well-annotated (WA) subset, instructing each model to output predictions for different phenotypic attributes, which generated 120,258 individual phenotype assignments. These assignments were then compared to the annotations provided by the bugphyzz database (**Supplementary Dataset 11**).

Assignment accuracy varied substantially across phenotypic attributes and models (Fig. 4A). Across all models, ‘spore formation’ was the most accurately assigned phenotype (mean balanced accuracy: 89.8%), while ‘biofilm formation’ proved most challenging (mean balanced accuracy: 53.0%), followed by ‘Gram staining’, which was largely attributable to poor performance on Gram-variable species. For example, while Gemini-2.5-pro performs best for spore formation, GPT-4.1-nano excels at cell shape assignment, with a balanced accuracy of 91.7%, substantially outperforming the other 30 models (Fig. 4B). Notably, it achieves very high performance on spirillum (100%), and tail (99.3%) morphologies. This primarily results from correctly classifying *Streptomyces* members as having tail-shaped morphology, while most other models misclassify them as bacillus-shaped. Overall, Gemini-2.5-pro achieved the top performance for two phenotypes, more than any other individual model.

**Figure 4:**
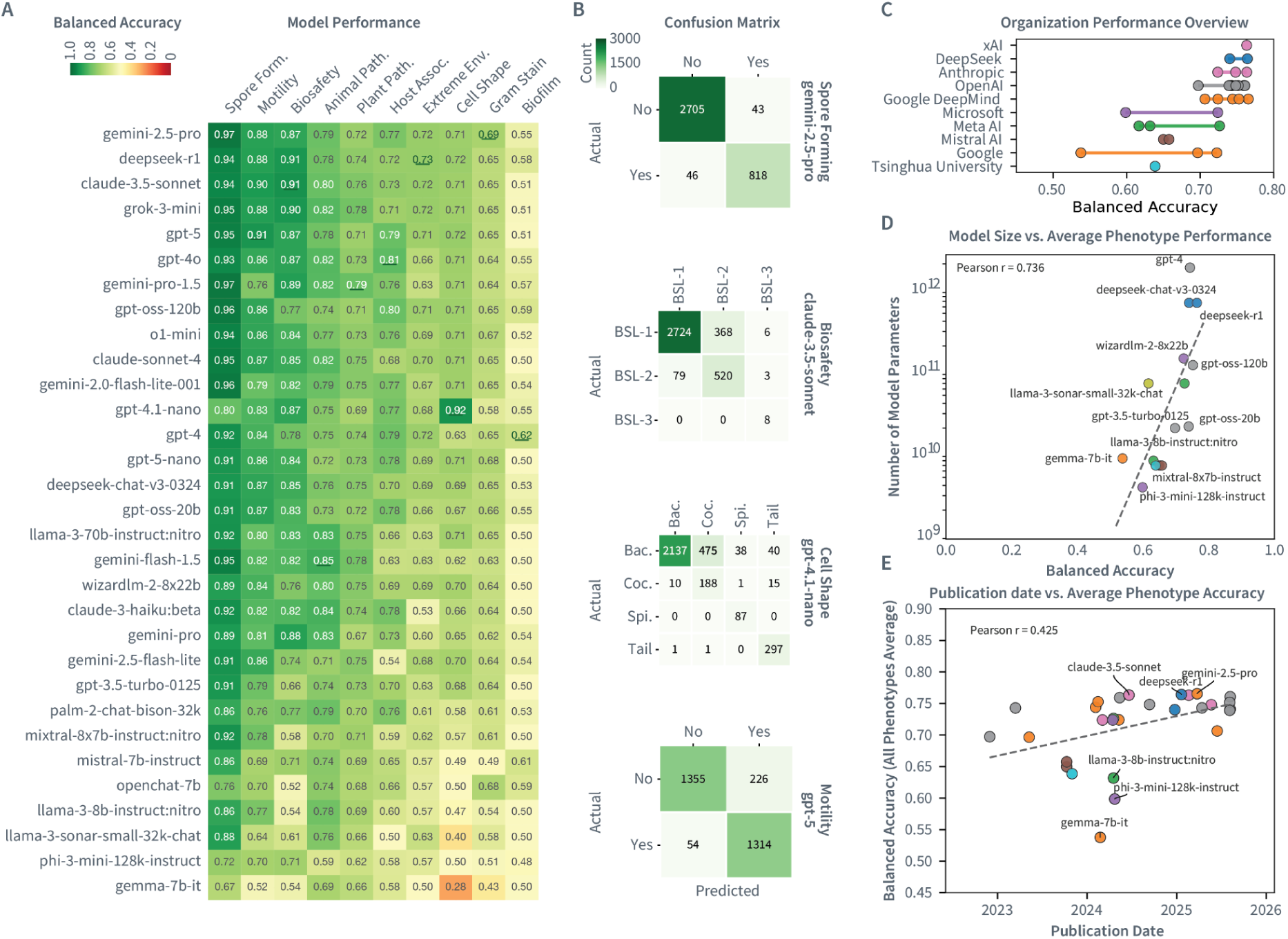
LLM performances in assigning phenotypes to microbial species. **A**) Balanced-accuracy heatmap for phenotype prediction across 31 models for 3876 species. Columns correspond to evaluated phenotypes; rows show models ordered by mean balanced accuracy. The best-performing model per phenotype is underlined. **B**) Confusion matrices for selected phenotype prediction tasks, showing model performance on specific traits. **C**) Horizontal ranges show organization-wise balanced accuracy. **D**) Relationship between model size and phenotype assignment accuracy. Balanced accuracy vs. parameter count (log scale), colored by organization, reveals a scaling trend (Pearson r=0.736). **E**) Publication date vs. accuracy indicates modest temporal gains (Pearson r=0.425).

We also evaluated the ability of LLMs to assign microbial biosafety levels (BSL-1, BSL-2, BSL-3), which correspond to increasing laboratory containment and safety requirements. The dataset is highly imbalanced, with far more BSL-1 classified species than species with higher biosafety levels (Fig. 1B). Biosafety level prediction represents the second most accurate predicted phenotype, with the best-performing model (claude-3.5-sonnet) achieving 91% balanced accuracy, outperforming even the newer claude-sonnet-4 (Fig. 4A, B). When aggregating performance by organization with at least two evaluated models (excluding xAI which had the model with highest accuracy), DeepSeek achieved the highest mean accuracy (75.2%, n=2 models), followed by Anthropic (74.5%, n=3), and OpenAI (74.4%, n=10). Notably, no single model dominated in performance across all phenotypes; different LLMs excelled at different prediction tasks (Fig. 4C). Notably, claude-3.5-sonnet correctly classified all eight BSL-3 species in the dataset. However, it also misclassified some BSL-1 species as BSL-3, specifically members of the *Brucella* genus (e.g., *B. cytisi*, *B. grignonensis*). Given the recent reclassification of *B. cytisi* from the genus *Ochrobactrum*, the model likely overestimated biosafety risk based on genetic proximity to other pathogenic *Brucella* species (Holzer et al. 2023). These misclassifications highlight how the benchmark reflects both the coverage and potential biases in training corpora, as well as genuine taxonomic ambiguities in the scientific literature.

We next assessed how phenotype classification performance scaled with model characteristics. Accuracy showed a strong positive correlation with model size (Pearson r=0.78; n=14 models with reported parameter counts) and a moderate correlation with publication date (Pearson r=0.32, n=29 models), suggesting that larger models substantially captured more phenotypic information in our evaluation) (Fig. 4D, E). The newest model releases (2024-2025) clustered with mean balanced accuracies above 0.76, whereas mid-sized models from 2023 typically scored 0.1-0.2 points lower. Together, these trends indicate both rapid recent progress in LLM capabilities and a pronounced model size effect, suggesting that accuracy will likely continue to improve with larger model architectures.

### Enhancing and extending phenotype annotations via LLM predictions

To further leverage high-accuracy predictions, we developed a strategy to identify the best-performing model for each phenotypic attribute. In this analysis, we used the same model for both knowledge group stratification and phenotype prediction, ensuring consistency in the evaluation process. We first examined the relationship between the LLM’s self-rated knowledge level (using query template 3) and its prediction performance (**Supplementary Dataset 9**). For analyzing knowledge-group trends, we applied stringent filtering criteria: we included only model-phenotype pairs with at least 30 labeled species within each knowledge group, and required each knowledge group to contribute at least 250 total observations across all models, before calculating group-level averages. Under these criteria, we observed a monotonic increase in balanced accuracy as self-rated knowledge level increased, consistent across models and phenotypes (Jonckheere–Terpstra trend test: z=5.02, p=2.54×10⁻⁷, Fig. 5A).

**Figure 5:**
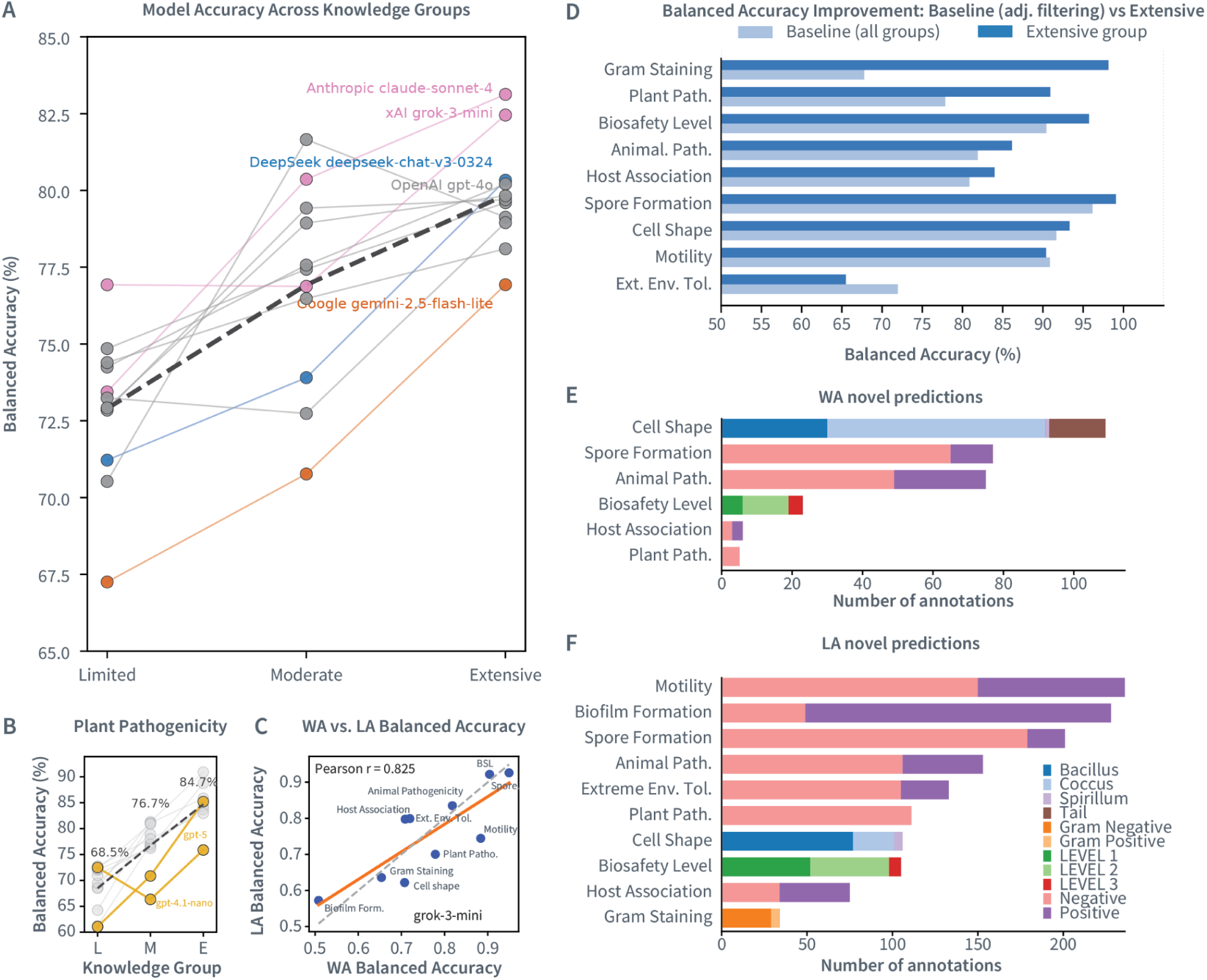
Improvement of microbial phenotype assignments with increasing knowledge-confidence of LLMs. **A)** Relationship between self-rated knowledge groups for models demonstrates consistent improvement with increased self-reported knowledge (mean Pearson r=0.75 across phenotypes; Jonckheere-Terpstra test p = 2.54 × 10⁻⁷). The dashed line represents the mean across models. **B**) Phenotype-specific knowledge scaling. Plant pathogenicity accuracy increased steadily from ‘Limited’ to ‘Extensive’ knowledge groups. The dashed line represents the mean. Yellow denotes results from OpenAI models. **C**) Cross-dataset performance consistency. Scatterplot shows the correlation between balanced accuracy on the WA and LA dataset for grok-3-mini across different phenotypes. Orange line represents linear fit, while dashed line represents the identity line (x=y). **D**) Performance gains from ‘Extensive’ knowledge filtering. Change in balanced accuracy when restricting to species self-rated as ‘Extensive’ knowledge versus all species, showing substantial improvements for eight of nine phenotypes. **E**) Novel high-confidence annotations generated by LLMs. Number of new phenotypic predictions for species lacking ground-truth data, for best performing model and cases where the model (**Supplementary Table 2**) self-rated its knowledge as ‘Extensive’ for each phenotype. **F**) Novel high-confidence annotations generated for the LA dataset. Number of new phenotypic predictions for species lacking ground-truth data in the LA set, using grok-3-mini and restricted to cases where the model self-rated its knowledge as ‘Extensive’.

This effect was especially pronounced for ‘plant pathogenicity’ (Benjamini-Hochberg adjusted p=1.53×10⁻⁷) and ‘Gram staining’ (BH-adjusted p=3.18×10⁻⁵), with all other traits except ‘extreme-environment tolerance’ remaining significant after false discovery rate control (Fig. 5B). Species self-rated with ‘Moderate’ or ‘Extensive’ knowledge groups averaged 76.2% and 79.7% balanced accuracy, respectively, compared with 72.6% in the ‘Limited’ knowledge group, demonstrating that self-assessed knowledge level serves as a useful indicator of prediction reliability. Consistent with this trend, within-phenotype correlations between knowledge level (called ordinally as ‘Limited’=1, ‘Moderate’=2, ‘Extensive’=3) and balanced accuracy were positive for eight of the nine phenotypes meeting our filtering criteria.

We generated a second dataset from bugphyzz, termed the Less Annotated (LA) set, which captures 15,256 species with sparse phenotype coverage and serves as a complementary benchmark to the curated WA subset. Overall, balanced accuracies on the LA and WA sets were highly correlated (Pearson r=0.825, Fig. 5C), suggesting that the relative strengths and weaknesses observed in the curated WA benchmark largely transferred to the sparser LA dataset. Accuracies remained high for traits like ‘spore formation’ (0.926) and ‘biosafety level’ (0.922). However, some phenotypes showed notable performance drops: ‘motility’ decreased from 0.884 (WA) to 0.744 (LA), and for ‘cell shape’, accuracy fell from 0.707 to 0.622. Balanced accuracy increased monotonically across knowledge groups: from 70.6% in the ‘Limited’ group to 77.5% in the ‘Moderate’ group (+6.9 percentage points), and reached 81.4% for the ‘Extensive’ group (+3.9 percentage points over ‘Moderate’). The monotonic gains across the three knowledge categories mirrors the pattern observed in the WA dataset, indicating that self-assessment reliability remains consistently even when applied to sparsely annotated species (**Supplementary Dataset 12**).

For each of the 10 phenotypes we compared the prediction accuracies of all LLMs within the ‘Extensive’ knowledge group. This approach identified optimal phenotype-model pairs, where certain models demonstrated superior performance for specific attributes. Restricting predictions to the ‘Extensive’ knowledge group yielded substantial accuracy gains for several phenotypes (Fig. 5D). ‘Gram staining’ showed the largest improvement (+28.7 points; from 69.5% to 98.1% balanced accuracy), followed by ‘plant pathogenicity’ (+11.5 points; from 79.4% to 90.9%).

‘Biosafety level’ and ‘host association’ each gained roughly 3-4 percentage points, whereas ‘motility’ and ‘extreme environment tolerance’ showed minimal change or slight decrease. These results demonstrate that filtering for high-confidence knowledge predictions can achieve very high predictive accuracies for many phenotype assignments.

Leveraging these optimal phenotype-model pairs (**Supplementary Table 3**), we extended annotations in the WA subset to fill gaps in the existing data. Using the best-performing model for each attribute, we generated 295 new high-confidence phenotype assignments across 256 species that previously lacked annotations for those traits (**Supplementary Dataset 13**). This targeted augmentation increased overall annotation completeness in the WA subset by 0.76 percentage-points, with the most substantial gains observed for ‘cell shape’ (+109 species annotated), ‘spore formation’ (+77), and ‘animal pathogenicity’ (+75). The LA evaluation set encompassed 15,256 species with sparse phenotype metadata (**Supplementary Table 1)**. Annotation completeness varied substantially by phenotype: while some traits such as ‘Gram staining’ were relatively complete (6.8% missing), others like ‘spore formation’ had substantial gaps, with over 20% of annotations absent. To test whether LLMs could accurately predict phenotypes despite this sparsity, we deployed grok-3-mini, which showed strong performance in the WA benchmark, to the entire LA dataset.

As the model’s self-rated knowledge scores correlate with prediction accuracy, we next applied a stringent filter to the grok-3-mini predictions: we retained only cases where (i) the model self-rated its knowledge as ‘Extensive’ for that species, and (ii) the LA dataset lacked any ground-truth annotation for that phenotype. This filtering yielded another 1,382 new, high-confidence phenotype assignments across 253 species (Fig. 5F**, Supplementary Dataset 14**), including 236 for ‘motility’ and 228 for ‘biofilm formation’, among others. While the confidence levels for these predictions may be lower than those on the WA subset, they provide valuable further metadata facilitating expert review and confirmation.

## Discussion

LLMs are increasingly used for integrating and synthesizing knowledge across diverse scientific domains, but they remain prone to generating plausible-sounding yet incorrect information, including clinically relevant hallucinations that can misguide organism identification and resistance profiles (Perkins et al. 2025). In this study, we systematically evaluated the strengths, limitations, and consistency of over 50 LLMs in deriving high-quality phenotype metadata for microbial species from texts. Microbial phenotypes underpin our understanding of microbial characteristics, functional roles, and applications, and are therefore a key to comprehensively and accurately describe in microbiology. Based on our results, this setup makes it possible to identify the conditions under which LLMs produce high-confidence, high-accuracy predictions and to focus downstream analyses on these reliable subsets. To enable reproducible and extensible evaluation, we developed an automated framework that uses standardized species sets, defined phenotype queries, and structured evaluation metrics and a self-assessment approach to systematically assess the propensity to hallucinate for different models. This framework facilitates continuous updates as new LLMs become available, providing a versioned benchmarking server with a browser interface that reports accuracy, calibration, and hallucination metrics and supports continuous re-evaluation as models evolve, with further applications for other data types in microbiology.

Application of LLMs in microbiology to date have targeted related problems with varying success, such as re-annotating sequencing records into environmental categories to classify microbial samples into ontology categories, including prediction of *E. coli* pathogen contamination risk from environmental metadata (Yoo and Rosen 2025). Other work evaluates extraction of biological terms (e.g., cell lines/genes) from NCBI/EBI BioSample text (Ikeda et al. 2025). Large-scale assessment on eukaryotic species using IUCN Red List information report strong knowledge retrieval, but rather weaker ecological reasoning (Uryu 2025). Recent successful applications of LLMs to biomedical data include using models with advanced retrieval strategies such as Retrieval-Augmented Generation (RAG), for example platforms like AskMicrobe that enrich general models with domain-specific context.

Our hallucination evaluation benchmark revealed clear performance stratification among models in their ability to recognize and correctly reject non-existent taxa. Generally, larger and more recent LLMs consistently outperformed smaller or early-generation models. The top performers, gpt-5-nano, gpt-5, and grok-4, showed markedly higher accuracy in identifying artificial binomial names as unknown. In contrast, smaller instruction-tuned models such as llama-3.2-3B-Instruct, Mistral-7B-Instruct, and older Gemma variants exhibited high hallucination rates on these data, often producing confident, yet incorrect classifications for synthetic taxa. Notably, even some widely deployed, large models such as gpt-4o demonstrated moderate hallucination tendencies, suggesting that model scale alone does not guarantee reliable uncertainty handling. Among open-weight models, gpt-oss-120b emerged as one of the best performers, demonstrating that proprietary systems do not uniformly outperform open alternatives by a large margin.

We next quantified how the amount of knowledge available on the internet for a given species relates to the LLM’s model confidence by measuring the model’s self-assessed knowledge level for real species and relating this to the number of Google search result counts for the same binomial names. Larger and more recent models generally showed better web-alignment, suggesting that both model scale and training recency improve knowledge calibration. However, we observed considerable variation even between models that are architecturally similar. When comparing knowledge-level self-assignments across models, we found varying degrees of agreement. As expected, well-characterized species showed high consensus across models. Interestingly, many low consensus cases could be traced to linguistic sources of confusion rather than genuine knowledge gaps: recent taxonomic reclassifications (*Metamycoplasma arthritidis*), Latin spelling variants (terra vs. terrae), concatenated epithets (*albiflaviniger*), or prefixes like “pseudo–” that cause spurious associations (*Rhodopseudomonas pseudopalustris*). These subtle variations in microbial nomenclature demonstrate that inter-model disagreement often reflects the inherent complexity and ambiguity of biological language rather than incomplete training.

Having evaluated how LLMs self-assess their knowledge and handle uncertainty, we examined how these capabilities translate to biological prediction tasks. Across the phenotype benchmark, no single model uniformly dominated performance, with most contemporary LLMs achieving similar overall accuracies (Fig. 4A). Performance varied systematically by phenotype: morphology-linked traits like spore formation, cell shape, and taxonomically stable characteristics such as biosafety level achieved consistently higher accuracy, whereas attributes like biofilm formation and Gram variability classification remained challenging. Several models, primarily older or smaller architectures, underperformed across multiple traits. Those patterns have important practical implications for model deployment. First, model selection should be phenotype-aware: selecting the best-performing model per trait can improve accuracy by up to 28 percentage points (e.g. Gram staining). Second, operational constraints such as inference costs and open-weight availability may influence model choice. Third, we observed a strong capacity effect, with accuracy correlated with both model size (Pearson r=0.78) and publication date (r=0.43), suggesting that both scale and training methodology continue to drive improvements.

Another key finding from our study is that the model’s self-assessed knowledge level about a species represents a reliable indicator of prediction accuracy, providing users with a systematic approach to distinguish between reliable and unreliable pieces of information. We saw a strong monotonic relationship between the LLM’s self-rated knowledge level and predictive performance across most phenotypic attributes, suggesting that LLMs are generally capable of assessing the confidence of their own outputs.

To demonstrate our findings and the utility of the resource, we applied this framework to prioritize novel phenotype assignments where no ground-truth data is present. This method combines high-confidence predictions for the well-annotated species set based on the best-performing model–phenotype combinations (tier 1), and high-confidence assignments for the model used on the extended less-annotated (LA) set (tier 2), delivering 1,382 novel, reliable phenotype annotations both for well-studied and understudied microorganisms. Principally, model confidence self-assessment can likely be applied to identify further types of high quality annotations for microbial taxa, such as properties of the environment they inhabit.

LLMs do not directly expose the source or evidence underlying each prediction, and we cannot determine whether relevant primary literature was included in the data. Given the opacity of most training corpora and the fact that many culture-based phenotypes are reported only in supplemental materials or specialized databases, further key evidence may currently still be underrepresented. Future improvements could further strengthen the outcome therefore. The phenotype selection and definition process could be made more structured, for example, by aligning trait descriptions with established ontologies such as ENVO/PATO or related phenotype vocabularies (Gkoutos et al. 2018; Buttigieg et al. 2013). Additionally, species-name-only prompting has inherent limitations for metagenomic lineages, which lack stable binomial names, recently reclassified taxa may be misidentified, and newly described species have minimal literature coverage, making zero-shot inference unreliable. Finally, our evaluation includes only general-purpose, publicly accessible LLMs; domain-specialized, fine-tuned biomedical models or agentic systems with retrieval augmentation may offer stronger phenotype coverage or better calibration.

Taken together, our results demonstrate that current LLMs encode substantial microbial knowledge and can accurately generate phenotype assignments with quantifiable reliability when paired with self-assessment filtering. LLMs thus provide a scalable, cost-efficient approach for accessing and synthesizing dispersed information from publicly available texts, though currently with varying performances for different phenotypes. As model capabilities continue to improve and domain-specific training becomes more feasible, LLM-based approaches combined with other data sources, such as microbial omics data, are likely to allow retrieving structured, high-quality microbiological metadata as a basis for integrative data analyses with AI-models and further experimental research.

## Supplementary Material

**Supplementary Dataset 1: Phenotypic and morphological attributes of microbial taxa used for LLM evaluation.** The table shows the ten phenotypic traits evaluated across two data subsets: the well-annotated (WA) subset with ≤5 missing values per species (n=3,876 species) and the less-annotated (LA) subset with sparse phenotypic coverage (n=15,256 species).

**Supplementary Dataset 2: Curated well-annotated (WA) microbial phenotype dataset derived from bugphyzz.** This table presents a refined dataset of microbial phenotypes, curated from the bugphyzz database, serving as the foundation for model comparison. The dataset encompasses a wide range of bacterial species and their associated phenotypic attributes.

**Supplementary Dataset 3: Curated less-annotated (LA) microbial phenotype dataset derived from bugphyzz.** This table presents a refined dataset of sparsely annotated microbial phenotypes, curated from the bugphyzz database, serving as the foundation for model comparison.

**Supplementary Dataset 4: Google search counts for binomial names in the WA subset.**

This table presents the Google search counts for species’ binomial names in the WA subset. The search counts were obtained using the Google API, reporting only exact matches for each binomial name. The table is sorted in descending order of search count.

**Supplementary Dataset 5: LLM query templates for knowledge group assessment and phenotype assignments**. This table presents query templates designed to assess an LLM’s self-rated knowledge about specific species and to generate JSON predictions of phenotypic attributes based on the organism’s binomial name.

**Supplementary Dataset 6: Artificial binomial names for LLM knowledge assessment validation.** This table presents 200 artificially generated binomial names used to evaluate the robustness of the LLMs’ confidence scoring system. Names are divided into four categories of 50 entries each: completely artificial names combining English words, pseudo-Latin names mimicking scientific nomenclature, hybrid names blending real and fabricated bacterial taxonomic elements, and other mixed constructs.

**Supplementary Dataset 7: LLM predictions for artificial binomial names.** This table presents the raw output from 57 different LLMs when presented with 200 artificial bacterial names using knowledge query template 3, our most detailed query template. The knowledge group assignments (NA, ‘Limited’, ‘Moderate’, or ‘Extensive’) represent the LLMs’ direct responses to these fabricated species names.

**Supplementary Dataset 8: LLM performance for artificial binomial names.** This table presents the hallucination rate (%) and hallucination-avoidance scores for 57 language models, each evaluated on 200 fabricated bacterial binomial names. Each response receives three points for an explicit refusal/NA, two for a ‘Limited’ knowledge claim, one for a ‘Moderate’ knowledge claim, and zero for ‘Extensive’ knowledge. The “Average Score” column reports the mean points per prompt, while the “Hallucination Rate” indicates the percentage of ‘Limited’, ‘Moderate’, or ‘Extensive’ answers.

**Supplementary Dataset 9:** Knowledge-level LLM predictions for the WA subset. This table presents the performance of 22 LLMs in predicting knowledge levels (Template Query 3).

**Supplementary Dataset 10: Shannon-entropy ranking of the benchmark species, computed from the mix of NA, limited, ‘Moderate’, and ‘Extensive’ self-reports supplied by the LLMs.** Entropy (in natural-log units) captures how evenly models distribute across the four self-knowledge bins. Values near zero reflect full consensus, whereas scores above ∼1.2 arise when comparable numbers of models select different labels, signaling ambiguous or hallucination-prone cases.

**Supplementary Dataset 11: Per-phenotype balanced accuracy for every language model evaluated on the WA dataset.** Each row lists the model name, phenotype field, and balanced accuracy derived from matched species predictions. Binary traits use the average of sensitivity and specificity, while multiclass phenotypes use macro recall (with the companion macro precision shown). Sample size reports how many species contributed to non-missing labels.

**Supplementary Dataset 12: Per-phenotype performance metrics for the LA dataset.** Only model–group–phenotype combinations with ≥30 paired observations are reported.

**Supplementary Dataset 13: Novel WA phenotype assignments.** Novel phenotype predictions spread across 256 unique species that were not covered by the ground-truth data before, using the best-performing phenotype–model combinations.

**Supplementary Dataset 14: Novel LA phenotype assignments.** Grok-3-mini and deepseek-chat-v3.1 phenotype calls paired with self-rated knowledge tiers for every LA species. Each row reports the species, knowledge group, phenotype, prediction, and ground-truth information if available.

## Acknowledgements

We acknowledge OpenAI for providing Research Credits, which were crucial for conducting this study. We utilized Anthropic’s Claude 4.5 Sonnet to assist with proofreading of this manuscript and OpenAI o1, OpenAI Codex and Claude Code for code generation for LLM-BioEval web page and the evaluation pipeline. The authors thoroughly reviewed and verified the accuracy of all code and content, maintaining full responsibility for the manuscript’s integrity.

## Author Contributions

P.C.M. developed the concept, implemented the evaluation pipeline and the code to produce all Figures, evaluated the models, and performed the statistical tests. N.S. contributed to the biological interpretation of the findings. G.R. contributed to the web server development. R.M. performed predictions using the deep learning model for comparison. Y.H. contributed to the ensemble model analysis. M.B., E.A.F., C.H., and A.C.M. provided guidance throughout the project. P.C.M., A.C.M., and N.S. wrote the manuscript, and all authors reviewed and approved the final version.

## Methods

### Online platform

The benchmarking platform provides real-time aggregated results through an online dashboard (https://llm-bioeval.bifo.helmholtz-hzi.de). Users can examine overall model performance, phenotype-specific metrics, and detailed error patterns. The platform is actively maintained, and newly released LLMs, both open-weight and API-based, are added on a continuous basis to keep the benchmark aligned with developments in the field. The portal provides programmatic access to the full dataset through JSON APIs and CSV exports, including endpoints for phenotype ground truth, model accuracy summaries, and raw prediction tables, and mirrors the database browser used internally so that users can inspect schema metadata and table sizes. We publish the complete SQLite database alongside these endpoints, allowing external groups to reproduce our analyses or integrate the annotations into new workflows. Because both services share the same data store, future model runs initiated via the administrative dashboard become visible on the public site automatically, ensuring that subsequent community contributions or newly released OpenRouter models can be incorporated without additional deployment steps.

### Data collection

Trait annotations were obtained from the harmonized BacDive tables distributed with the bugphyzz R package. Using dplyr and stringr, each phenotype table was filtered to species-rank records with binomial Latin names (two tokens, excluding “sp.” designations) and to annotations reported with frequency “always” or “usually.” The workflow retained 13 phenotypes—motility, gram stain, aerophilicity, extreme environment adaptation, biofilm formation, animal pathogen status, biosafety level, health association, host association, plant pathogenicity, spore formation, hemolysis, and cell shape. For hemolysis we required alpha, beta, or gamma designations, and cell-shape categories were limited to those occurring in at least 1% of observations. Within each trait table, duplicate species entries were collapsed to a single record before full-joining all traits by NCBI taxonomy identifier and species name. We flagged species annotated for more than 60% of traits as “highly annotated.” The resulting table comprises 19 066 species; trait coverage ranges from 95% (gram stain) and 82% (spore formation) to 57% (aerophilicity) and 28% (motility).

### Administrative Inference Dashboard

We developed a local administrative dashboard to coordinate high-throughput inference with the pipeline. The Flask and Socket.IO application presents a browser-based control surface that couples curated species lists with paired system and user templates, selects the target language model, and records all job metadata in a unified SQLite backend. Inference calls are issued through the OpenRouter API with deterministic decoding (temperature fixed at 0.0), ensuring run-to-run reproducibility and full provenance for every prediction. The admin interface is model-agnostic and can launch inference against any OpenRouter-registered model (currently 300+), so the same controls apply across proprietary, open, or community releases without code changes.

Job execution is orchestrated by a processing manager that manages a configurable thread pool (default 10 concurrent workers) and rate limits, while streaming real-time status updates and logs to the client via WebSocket events. Complementary REST endpoints expose the same controls, job creation, pause/resume/stop operations, and dynamic adjustment of concurrency or request rate, allowing automated monitoring pipelines to intervene without manual interaction. Each inference retains its raw request and response payloads, parsed phenotype outputs, timestamps, and error diagnostics, enabling post hoc auditing or reanalysis. Inference requires an OpenRouter API key exported to the runtime environment, so every job is authenticated against the same provider configuration before dispatch. Completed predictions and their validation metadata are archived in the unified SQLite database.

Template management and validation are integral to the dashboard. System and user prompt templates are versioned within the interface, species lists are curated and linked to jobs, and optional benchmark datasets provide ground-truth phenotype labels. During execution, responses are parsed into the predefined schema, written to the unified database, and scored immediately when benchmarks are present. The platform can rerun failed species, reparse historical predictions after template updates, or replay entire jobs, providing a reproducible, queryable record that underpins downstream analyses reported in the manuscript.

Validation of predictions is governed by a parallel set of templates that define the expected JSON schema and permissible vocabularies for each phenotype. For every job, the dashboard stores both the inference prompt pair and the matching validation template, ensuring that returned fields are parsed and normalized consistently. When benchmark datasets are provided, the processing manager compares each model prediction against the ground-truth phenotypes in real time, logs concordant and discordant cases, and flags conflicts for analyst review before downstream export. This workflow lets us correct template mismatches rapidly, rerun only the affected species to keep inference costs down.

### Synthetic calibration benchmark

We queried each LLM with a dedicated set of knowledge-assessment prompts that return a single field knowledge_group. Three prompt variants were authored (templates 1–3), increasing the descriptive context supplied to the model. Template 1 provides minimal category definitions (‘Limited’, ‘Moderate’, ‘Extensive’), template 2 elaborates with qualitative cues, and template 3 enumerates quantitative heuristics (e.g., approximate literature counts, culture availability) and explicitly instructs the model to answer “NA” whenever the species is unrecognized or insufficiently documented. All prompts are executed through the administrative inference stack at temperature 0.0, validated with the shared PredictionValidator, and any free-form responses are normalized to the canonical vocabulary; synonyms such as “minimal,” “medium,” or “high” are mapped to ‘Limited’, ‘Moderate’, and ‘Extensive’, while unparsable or empty payloads are flagged as no_result and transport errors as inference_failed.

To probe hallucination behaviour, we assembled a benchmark of 200 artificial binomial names. The set contains four equally sized categories (n = 50 each): English word pairs, purely Latin-like inventions, hybrids pairing a real genus with an invented epithet, and the converse (invented genus with a real species epithet). Each entry retains its category label so results can be stratified by name realism. The same species list is issued to every model across the three knowledge templates when we study prompt verbosity effects, and the full panel of 57 models is evaluated with template 3 to quantify NA compliance.

All models are accessed through OpenRouter via the administrative dashboard, yielding 11 400 knowledge judgements (57 models × 200 synthetic names) for the primary hallucination analysis plus additional runs for the lower-verbosity templates on a representative subset of models (e.g., GPT-4, Claude 3.5 Haiku, Llama 3 8B). For each model we tabulate the counts of NA, ‘Limited’, ‘Moderate’, ‘Extensive’, no_result, and inference_failed outcomes; limited/moderate/extensive answers on an artificial species are labelled hallucinations, NA counts as a correct rejection, and the latter two categories track non-responsive failures. Model-level hallucination rates are computed as (‘Limited’ +‘Moderate’+‘Extensive’) / total, and we report both per-model percentages and the aggregate distribution. The same normalised outputs feed downstream agreement analyses, Shannon entropy across models, disagreement heatmaps, and quality score tables, that are rendered through the knowledge-calibration research components.

### Ground-truth benchmark sets

Harmonized annotations (19 066 species across 13 phenotypes) generated by the bugphyzz pipeline were imported into the administrative dashboard, where trait names and vocabularies were standardized against the internal schema and stored in the unified SQLite database. Although the full trait panel is preserved for provenance, benchmark analyses are restricted to the ten phenotypes that exhibited single-valued annotations (one categorical state per species) and passed our predefined coverage threshold; phenotypes permitting simultaneous categories or falling below coverage were omitted to ensure interpretability of accuracy and calibration estimates. Each inference job therefore operates against the same canonical reference table, allowing predictions to be validated automatically without manual data handling and guaranteeing that downstream performance metrics reflect a consistent, reproducible ground-truth baseline.

### Knowledge calibration on real species

We evaluated model concordance on the well-annotated (WA) benchmark list of 3 885 bacterial species, which pairs each binomial name with a Google search-count proxy for literature prevalence. Every model in the panel was run through the knowledge-level prompt family, using template 3 for the primary analysis so that predictions were constrained to the vocabulary {limited, ‘Moderate’, ‘Extensive’, NA}. Inference calls were made via the pipeline (temperature 0.0) and validated with the shared PredictionValidator, which maps synonymous responses (for example “minimal,” “high,” “unknown”) onto the canonical categories and records empty parses as no_result and transport errors as inference_failed. The resulting classifications are stored in the unified SQLite database together with the originating template identifiers, enabling downstream aggregation without repeated API calls.

Species-level agreement summaries were generated directly from the database-backed API. For each species we collected all completed template 3 predictions across models, required at least five model outputs to retain the species, and tabulated the distribution of NA/‘Limited’/‘Moderate’/‘Extensive’ labels. Shannon entropy (H=-\sum_i p_i \log p_i) on the normalized counts serves as our measure of disagreement, with (H=0) indicating unanimous agreement. We also track the number of distinct knowledge groups assigned to each species and the majority-class fraction to distinguish total consensus (entropy ≈ 0, single class) from partial splits across two, three, or four categories. These statistics populate the entropy-ranking tables, disagreement heatmaps, and the counts cited in the Results (e.g., 91 unanimous species, 1 046 split across two groups). The same aggregation produces per-model distributions—number of species labelled NA, ‘Limited’, etc.—used for the performer rankings and stacked-bar visualizations.

To relate model confidence to real-world information availability, we aligned the knowledge assignments against the google search counts bundled in the WA file. An API endpoint joins the inference results with the search-count column, filters to completed template 3 predictions, and converts categorical knowledge levels to ordinal scores (‘Limited’= 1, ‘Moderate’= 2, ‘Extensive’= 3). NA classifications are retained for reporting but excluded from the correlation itself. We apply a log₁₀ transformation to search counts to dampen the heavy-tailed distribution and compute model-specific Pearson correlations between the transformed counts and knowledge scores, along with summary statistics by knowledge level (mean and s.d. of log-hit counts). These correlations, the “web alignment” scores reported are accompanied by the underlying knowledge distributions, revealing, for example, that models with high correlation often allocate more species to NA and fewer to ‘Extensive’, whereas over-confident models show inflated ‘Extensive’ counts and weaker alignment with public information proxies.

### Phenotype Benchmark

Phenotype evaluation draws on two curated species panels. The well-annotated (WA) benchmark contains 3,885 species and Google search-count proxies for literature prevalence. A complementary low-annotation (LA) list covers 15,256 species to probe generalization beyond the high-coverage subset. Both datasets include the 10 single-valued traits used throughout the study, gram staining, motility, extreme-environment tolerance, biofilm formation, animal pathogenicity, biosafety level, host association, plant pathogenicity, spore formation, and cell shape, while traits that admit multiple simultaneous categories or suffer from sparse labels (e.g., aerophilicity, health association, hemolysis) are excluded to keep accuracy estimates better interpretable.

All phenotype assignments are generated with the deterministic inference pipeline described above: a structured phenotype prompt elicits JSON-formatted responses that are normalized to the canonical vocabularies. For each model–phenotype pair we align predictions with the ground-truth labels, discard entries where either side is missing, and compute balanced accuracy. Binary traits use the mean of sensitivity and specificity; multiclass traits use macro-averaged recall. These metrics feed the accuracy heatmaps, per-phenotype leaderboards, and confusion matrices reported in the Results, and the same scoring routine is applied when evaluating the LA cohort or filtered subsets of species.

To assess how self-declared knowledge levels modulate performance, we join each phenotype assignment with the corresponding knowledge classification that the same model produced for the same species. Accuracy is recomputed within the ‘Limited’, ‘Moderate’, and ‘Extensive’ knowledge strata, provided at least 30 paired observations per phenotype and 250 samples per knowledge group are available. Responses are normalized so that synonyms (e.g., “minimal,” “medium,” “high”) map to the canonical categories, while NA entries are tracked but excluded from the accuracy aggregates. We summarize these conditioned accuracies for the sample-rich models, plot per-phenotype trends, and apply Jonckheere–Terpstra tests with Benjamini–Hochberg correction to evaluate whether performance increases monotonically from limited to ‘Extensive’ knowledge. Companion analyses compare baseline (unfiltered) scores with ‘Extensive’-only subsets and measure how knowledge groups explain variance in phenotype accuracy.

Model-level averages are further related to architecture and provenance. We require that a model contribute at least 100 scored samples per phenotype before including it in the aggregate “all-phenotype” balanced accuracy. Parameter counts, release years, and provider labels are drawn from a curated metadata table, enabling correlation of log₁₀ parameter count with average balanced accuracy, temporal trend analyses across release cohorts, and organisation-level comparisons. Linear fits and Pearson correlations quantify these relationships, while color-coded scatter plots highlight differences between providers. Publication date and parameter count of models were sourced from Epoch.ai (Accessed September 2025)

