## Supplemental Information for "Comparative Assessment of Large Language Models for Microbial Phenotype Annotation"

### Supplementary information

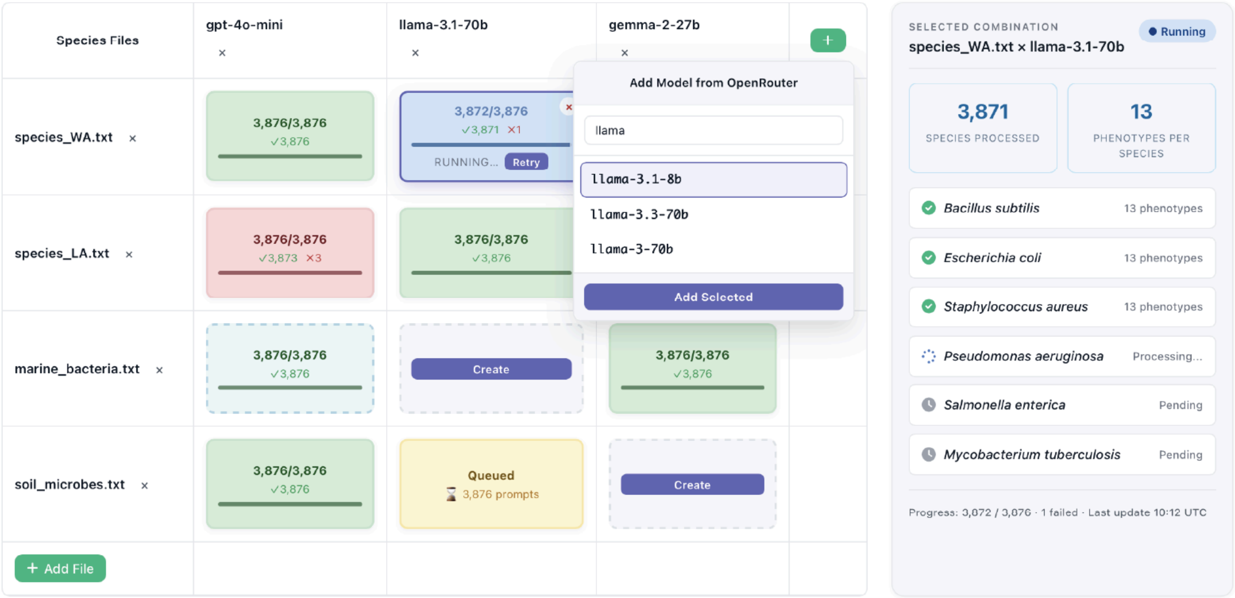


**Supplementary Figure 1: Admin Interface Overview.** The pipeline provides a browser-based administrative interface for orchestrating large-scale phenotype assignment experiments. The interface enables systematic evaluation of multiple LLM models across diverse species collections, with real-time progress monitoring and automated job scheduling. Users can dynamically expand the model roster through OpenRouter integration (dropdown shown), track prediction status for each species-model combination, and access detailed extraction statistics including species processed, success rates, and failure counts for individual jobs.

**Table S1**: Phenotypic and morphological attributes of microbial taxa used for LLM evaluation.The table shows the ten phenotypic traits evaluated across two data subsets: the well-annotated (WA) subset with ≤5 missing values per species (n=3876 species) and the less-annotated (LA) subset with sparse phenotypic coverage (n=15,256 species).

| **Label** | **Type** | **Targets** | **Annotated fraction [in %]** | |
|---|---|---|---|---|
| | | | **WA subset** | **LA subset** |
| **Motility** | Binary | `[TRUE, FALSE]` | 76.8 | 42.8 |
| **Spore formation** | Binary | `[TRUE, FALSE]` | 94 | 78.4 |
| **Gram staining** | Multiclass | `[gram stain negative, gram stain positive, gram stain variable]` | 99.8 | 93.2 |

| | | | | |
|---|---|---|---|---|
| **Cell shape** | Multiclass | `[bacillus, coccus, spirillum, tail]` | 85.6 | 32.6 |
| **Host Association** | Binary | `[TRUE, FALSE]` | 69.2 | 11.4 |
| **Plant pathogenicity** | Binary | `[TRUE, FALSE]` | 97.0 | 73.0 |
| **Biosafety level** | Multiclass | `[biosafety level 1, biosafety level 2, biosafety level 3]` | 96.5 | 84.2 |
| **Extreme environment tolerance** | Binary | `[TRUE, FALSE]` | 87.1 | 17.5 |
| **Animal Pathogenicity** | Binary | `[TRUE, FALSE]` | 53.2 | 7.3 |
| **Biofilm formation** | Binary | `[TRUE, FALSE]` | 6.6 | 1.0 |

**Table S2: Best-performing language models selected for individual microbial phenotype assignments.** Each phenotype is paired with the language model achieving the highest balanced accuracy on species in the “Extensive” knowledge group within the WA dataset.

| Phenotype | Balanced accuracy | Best model (used for inference and knowledge grouping) | Sample size |
|---|---|---|---|
| Spore Formation | 96.6% | Google gemini-2.5-pro | 3,612 |
| Cell Shape | 91.7% | Openai gpt-4.1-nano | 3,290 |
| Biosafety Level | 91.4% | Anthropic claude-3.5-sonnet | 3,708 |
| Motility | 90.9% | Openai gpt-5 | 2,949 |
| Animal Pathogenicity | 84.9% | Google gemini-flash-1.5 | 2,050 |
| Host Association | 80.8% | Openai gpt-4o | 2,665 |
| Plant Pathogenicity | 79.4% | Google gemini-pro-1.5 | 3,723 |
| Extreme Environment Tolerance | 72.5% | Deepseek deepseek-r1 | 3,350 |
| Gram Staining | 69.5% | Google gemini-2.5-pro | 3,840 |
| Biofilm Formation | 61.7% | Openai gpt-4 | 253 |

**Table S3**: **Best-performing model selection for phenotype prediction using high-confidence filtering.** Best-performing model for each phenotype for species self-rated as having “Extensive” knowledge. Phenotypes where this filtering reduces accuracy (Motility, Extreme environment tolerance) are excluded.

| Phenotype | Model | Balanced Accuracy (%) | Samples |
|---|---|---|---|
| Gram Staining | xAI grok-3-mini | 98.14 | 402 |
| Animal Pathogenicity | xAI grok-3-mini | 86.14 | 327 |
| Biosafety Level | xAI grok-3-mini | 95.72 | 379 |
| Host Association | Anthropic claude-sonnet-4 | 83.99 | 259 |
| Plant Pathogenicity | Anthropic claude-sonnet-4 | 90.91 | 260 |
| Spore Formation | OpenAI gpt-oss-120b | 99.07 | 351 |
| Cell Shape | OpenAI gpt-4.1-nano | 93.31 | 658 |
